# Career incentives make Registered Reports unattractive under realistic academic conditions

**DOI:** 10.64898/2026.07.17.739248

**Authors:** Anne M. Scheel, Leo Tiokhin, Daniël Lakens

**Affiliations:** Utrecht University; Deciding Better; Eindhoven University of Technology

## Abstract

Registered Reports are a publication format designed to reduce publication bias by guaranteeing publication before results are known. This guarantee is considered attractive in the prevailing ‘publish or perish’ culture. But would researchers always prefer such a low-risk option? When strong results allow publication in prestigious journals, the results-dependent nature of standard publications can actually be an advantage for authors. Given this trade-off, the choice between publication formats represents decision making under risk, and researchers may not always be risk-averse. To better understand under which circumstances Registered Reports are likely to be popular, we draw on risk-sensitivity theory from behavioural ecology and develop an agent-based model that explores how four factors shape publication strategies: whether additional publications yield diminishing or increasing career returns, the pace at which studies are completed, publication targets, and competition. We find that many realistic configurations lead researchers to avoid Registered Reports, even when Registered Reports are nearly as valuable as the best possible standard publication. Intense competition and low empirical pace can make the format unviable, while high empirical pace attenuates most other effects. Given prevailing career incentives in many fields, these results suggest that the market share of Registered Reports may remain trivially low without additional incentives or interventions.

---

Registered Reports are an article format designed to reduce publication bias and ‘questionable research practices’ (QRPs), which distort the published record of research findings in many scientific disciplines (Chambers, 2013; Franco, Malhotra, & Simonovits, 2014; Fraser, Parker, Nakagawa, Barnett, & Fidler, 2018; Gopalakrishna et al., 2022; John, Loewenstein, & Prelec, 2012). In this format, peer review takes place before data collection and the decision to publish is made before authors, reviewers, and editors know the study results. In addition to preventing editors from selectively rejecting unfavourable results (in particular negative or null results), this is thought to remove incentives for authors to hide, embellish, or misrepresent results because publication no longer depends on them (Chambers, Dienes, McIntosh, Rotshtein, & Willmes, 2015). Initial evidence from psychology and neighbouring disciplines shows that Registered Reports indeed contain much higher rates of negative results than the standard literature (Allen & Mehler, 2019; O’Mahony, 2023; Scheel, Schijen, & Lakens, 2021; Toth et al., 2021).

Advocates of the format have argued that the pre-data publication guarantee should make Registered Reports particularly attractive to researchers (e.g., Chambers & Tzavella, 2022) because it reduces uncertainty about whether and where a study will be published before authors have invested in conducting it. This provides relative safety in a research climate that involves substantial publication pressure (Gopalakrishna et al., 2022; Miller, Taylor, & Bedeian, 2011; Tijdink, Vergouwen, & Smulders, 2013; van Dalen & Henkens, 2012). The argument assumes that researchers’ choices are shaped by strategic concerns about publishability — but if they are, it is unlikely that such concerns would always cause risk aversion. Instead, researchers’ willingness to take risks likely varies as a function of available resources, time pressure, competition, and other factors. As a consequence, Registered Reports might remain unpopular in certain situations without additional incentives or interventions. And indeed, although the uptake of Registered Reports has grown rapidly (Chambers & Tzavella, 2022), their market share is much smaller than expected if scientists saw Registered Reports as unreservedly beneficial for their careers. Here, we examine under which circumstances Registered Reports remain an unattractive publication format with an agent-based simulation, in which we model authors’ choices between publication formats as decision making under risk.

## Design and function of Registered Reports

The review process of Registered Reports is split into two stages. At Stage 1, reviewers evaluate a pre-study protocol containing the research questions, hypotheses, methods, and analysis plan. In case of a positive decision, the journal issues an ‘in-principle acceptance’ and commits to publishing the eventual report regardless of the eventual results. Authors start data collection after in-principle acceptance. At Stage 2, the final report is peer-reviewed a second time to verify adherence to the protocol, data quality, and whether conclusions are justified by the evidence. Manuscripts can still be rejected at Stage 2, but only for substantial violations of the protocol or uninformative data (e.g., caused by equipment failure).

Through this process, Registered Reports address selective reporting and exploitation of flexibility during data analysis, two practices that bias scientific evidence and contribute to poor replicability and research waste (Chalmers & Glasziou, 2009; Ferguson & Heene, 2012; Wagenmakers, Wetzels, Borsboom, van der Maas, & Kievit, 2012). Selective reporting can occur at the level of whole studies — when editors and reviewers disproportionately reject submissions with negative results (‘reviewer bias,’ Atkinson, Furlong, & Wampold, 1982; Mahoney, 1977), or when researchers fail to submit studies with negative results for publication (‘file-drawering,’ Ensinck & Lakens, 2025; Franco et al., 2014; Rosenthal, 1979) — but also at the level of individual hypotheses or analyses, when researchers report only those that yielded favourable results (Gopalakrishna et al., 2022; John et al., 2012; Simmons, Nelson, & Simonsohn, 2011). Registered Reports’ in-principle acceptance issued at Stage 1 addresses study-level selective reporting: Editors and reviewers cannot reject the Stage-2 report based on the results, which also reduces the incentives for authors to file-drawer negative outcomes. The Stage-1 protocol additionally acts as a preregistration that constrains hypothesis-and analysis-level flexibility, with reviewers tasked to flag any undisclosed deviations from it during Stage-2 review.

Registered Reports were first launched in 2013 at the journal *Cortex* (Chambers, 2013) and are now offered by over 300 journals. Empirically, they appear to work as intended: Registered Reports show markedly lower rates of positive results than standard reports (Allen & Mehler, 2019; O’Mahony, 2023; Scheel et al., 2021; Toth et al., 2021). A comparison with matched controls found that they have larger sample sizes and higher methodological quality (Soderberg et al., 2021), suggesting that the lower positive result rate is not an artefact of reduced statistical power.

## Author incentives for Registered Reports

Registered Reports are generally thought to ‘[neutralise] bad incentives’ (Chambers, 2013, p. 609): Once authors have committed to the format, they can improve their publication chances only via the proposed research question and methods, and results are no longer a target to ‘hack’ or select on.^1^ The incentives for choosing the Registered Reports route in the first place, however, are less clear. Advocates of the format have argued that Registered Reports ‘serve the interests of individual scientists’ (p. 12, Chambers & Tzavella, 2022) because the format reduces the risk of investing in research projects whose results turn out to be difficult to publish: Under publication pressure, the pre-data publication guarantee removes what Chambers has called the ‘*p*-value lottery’ (Chambers, 2021) of standard publications. This argument assumes that researchers a) are under pressure to amass journal publications (which remain a central currency for hiring and promotion decisions, R. Müller, 2014; van Dalen & Henkens, 2012) and b) face shortfalls in publication output when their studies yield negative results.

These two assumptions are plausible, but publication decisions are influenced by additional factors. Typically, authors care not only about their studies being published, but also about the reputation of the publishing journal and subsequent citation rates (which are causally influenced by journal rank, Traag, 2021). High-impact journals appear to select more strongly on results (Ceausu et al., 2018; Siontis, Evangelou, & Ioannidis, 2011). Since Registered Reports are selected for publication before results are known, it may be no accident that very few high-impact journals currently offer the format.^2^ This means that although Registered Reports protect authors from publication failure, they also limit the maximum payoff that a study could provide. In other words, the *range* of possible outcomes is smaller than in standard reports.

In standard reports, career-relevant payoffs per study vary from very low — when authors file-drawer a manuscript because the publication chances do not justify the costs (Ensinck & Lakens, 2025) — to very high, when a manuscript is published in a high-impact journal and frequently cited. Compared to this, Registered Reports reduce variance at the lower end of the range by minimising the chances of no publication at all, but also at the upper end: Upon receiving in-principle acceptance, most authors know that their study will not be published in a high-impact journal. For standard-report authors, in contrast, the chance of a high-impact publication persists — at least hypothetically — until the results are in. Therefore, as long as the payoff associated with a published Registered Report is not always on par with the best possible outcome of the standard route, there will be situations in which gambling on a high-impact standard publication is more beneficial for researchers.

## Publication strategies as decision making under risk

Because Registered Reports and the standard publication route differ in payoff variance, authors’ choice between them represents *decision making under risk*. Here, we draw on risk-sensitivity theory from behavioural ecology to model factors that influence risk preferences and simulate their effects on researchers’ publication strategies. Following Winterhalder, Lu, & Tucker (1999), we define *risk* as ‘unpredictable variation in the outcome of a behavior, with consequences for an organism’s fitness or utility’ (p. 302).

*Risk aversion* thus means preferring a low-variance option over a high-variance option, and *risk proneness* the reverse. Organisms are *risk sensitive* when they are sensitive not only to the average of outcomes of different options, but also to their variance.

Risk-sensitivity theory is a normative theory originally developed to determine optimal foraging strategies for animals choosing between low-variance and high-variance food sources (Kacelnik & Bateson, 1996). Organisms are predicted to be sensitive to such differences when payoffs have non-linear consequences for survival or reproductive fitness, for example when additional food yields diminishing returns or when amounts below a threshold cause starvation. The predictions of risk-sensitivity theory overlap substantially with those of expected utility theory and prospect theory, but the central currency is fitness instead of utility. Despite its initially narrow scope, risk-sensitivity theory has proven itself as a powerful framework for explaining risk-sensitive behaviour across a wide range of situations and species, including humans (Mishra, 2014; Winterhalder et al., 1999).

## The present study

We apply risk-sensitivity theory to the situation of researchers choosing between Registered Reports and the standard publication route. Using a simulation model, we explore how four risk-relevant aspects of academic careers affect publication strategies: (1) whether additional publications yield linear, decreasing, or increasing returns for career success, (2) empirical pace (the frequency at which studies can be completed), (3) publication targets that must be met to continue or further one’s career, and (4) competition. Our goal is to understand in which circumstances Registered Reports should be particularly attractive or unattractive for researchers. The results of this analysis may help anticipate research fields and career stages in which the format is unlikely to take foot without additional changes to norms, incentives, or policy.

## Conceptual application of risk-sensitivity theory to publication decisions

In the context of academic careers, risk-sensitivity theory’s focus on reproductive fitness as the central outcome may be seen as misguided. But as argued by Smaldino & McElreath (2016), academic research satisfies the three requirements for natural selection: variation (e.g., in research practices), consequences of this variation for survival and reproduction (e.g., some practices increase the chances of staying in academia and of being copied by others), and heritability (e.g., PhD advisors passing their research practices on to their students). Analogous to natural selection, a competitive job market creates bottlenecks that select for specific behaviours. In academia, these are behaviours which (whether or not intentionally) increase a researcher’s chances of achieving tenure (Smaldino & McElreath, 2016; for a similar approach, see also Higginson & Munafò, 2016). We therefore conceptualise fitness as career success. This does not imply that career success is the only or the proximal motivation for researchers’ behaviour, just as evolutionary theory does not imply that reproductive success is the only motivation for human behaviour. However, we do assume that selection for career-promoting behaviours has a noticeable impact on research practice.

### Non-linear fitness functions

The first and most ubiquitous factor that leads individuals to be risk sensitive is a non-linear relationship between behavioural outcomes (e.g., harvested food items, publications) and fitness (Kacelnik & Bateson, 1997). Consider two options, *O_safe_* and *O_risky_*. *O_safe_* always provides the same payoff *b_safe_*, whereas *O_risky_* gives either a low payoff *b_low_* or a high payoff *b_high_*, each with probability 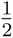 . When 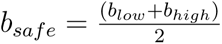, *O_safe_* and *O_risky_* have the same expected payoff. However, an individual should be indifferent between the two options only if the consequences of their payoffs for fitness are linear. As illustrated in Figure 1, *O_safe_* provides greater expected fitness than *O_risky_* when the fitness function is concave (diminishing returns), but lower expected fitness when the function is convex (increasing returns). Preferences should thus shift towards risk aversion when returns are diminishing and towards risk proneness when returns are increasing.

**Figure 1.**
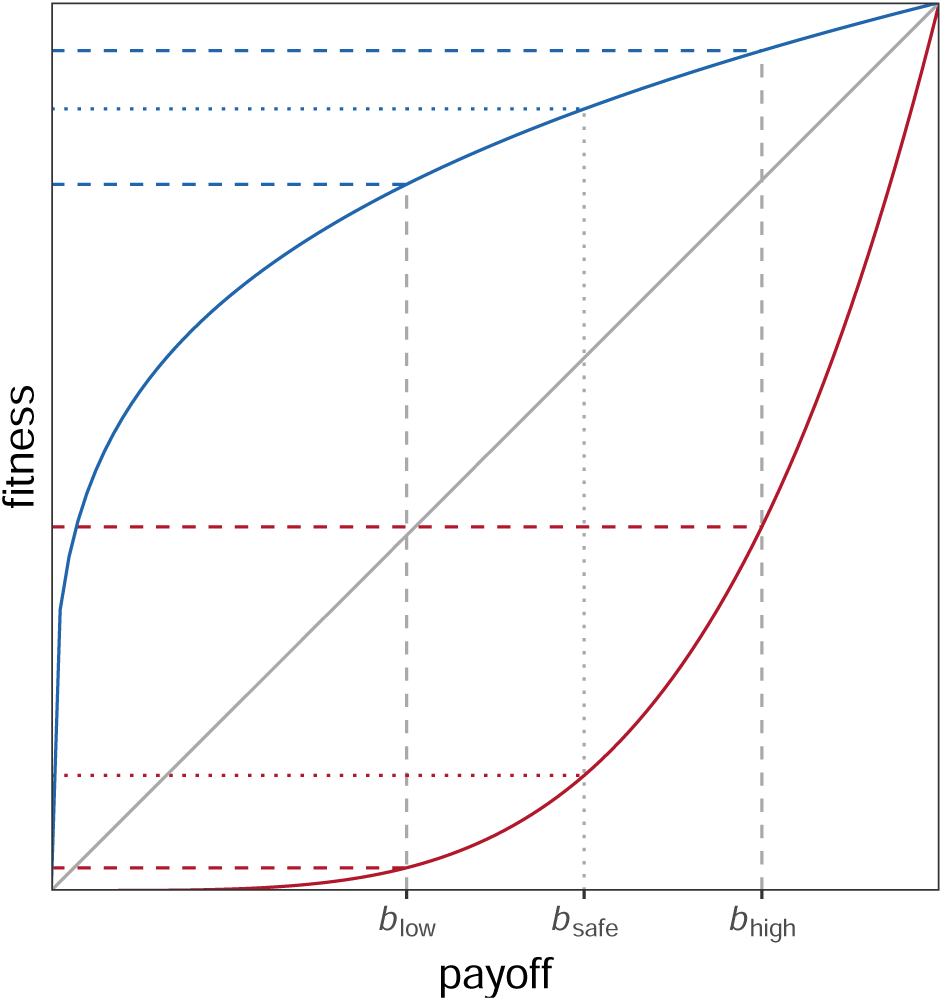
Consequences of non-linear fitness functions. Payoffs *b_low_*, *b_safe_*, and *b_high_* are converted into fitness with a diminishing (blue), linear (grey), or increasing (red) returns function.

Non-linear relationships are arguably the norm in the natural world and linear relationships the exception. This holds for academia as well: Any additional publication may have an accelerated effect on career success for early-career researchers, but will seem negligible for established professors.

### Number of decision events before evaluation

A second risk-relevant factor is the number of decision events taking place before an individual’s fitness is evaluated. When a risky option is chosen repeatedly, the average of accumulating payoffs approaches the long-run expected payoff. The risky option then becomes relatively more attractive because the danger of only ever acquiring the lowest possible payoff diminishes. However, this only holds for repeated decisions *before* fitness is evaluated: When fitness is evaluated after a single decision event, extreme outcomes leading to zero fitness (i.e., death or an ultimate failure to reproduce) are more likely and risk-averse strategies may be more adaptive, even when risk-prone strategies produce better outcomes within any one generation (Haaland, Wright, & Ratikainen, 2019).^3^

For the purpose of the present study, ‘decision events’ refer to researchers’ choices of whether to conduct a Registered Report or pursue the standard publication route. The number of decision events before evaluation reflects the number of empirical projects that a researcher can complete before their publication record is considered for hiring, promotion, or grant funding decisions. We call this parameter ‘empirical pace’. Empirical pace may differ between research areas that vary in the speed and cost of data collection (e.g., online questionnaires *vs* fMRI studies), between labs that vary in funding and manpower, and even between career stages, as seniority often comes with increased resources (R. Müller, 2014), while junior researchers face short contracts that limit the time for producing output before their next job application.

### Survival thresholds and competition

A final important factor for risk-sensitive behaviour are thresholds for survival and reproduction (Hurly, 2003; Winterhalder et al., 1999). Survival thresholds are cutoff points below which an individual’s fitness drops to zero, for example due to starvation. Individuals should be risk-averse when a low-variance option reliably meets the threshold, and risk-prone when only a higher-variance option offers a chance of reaching it (Figure 2).

**Figure 2.**
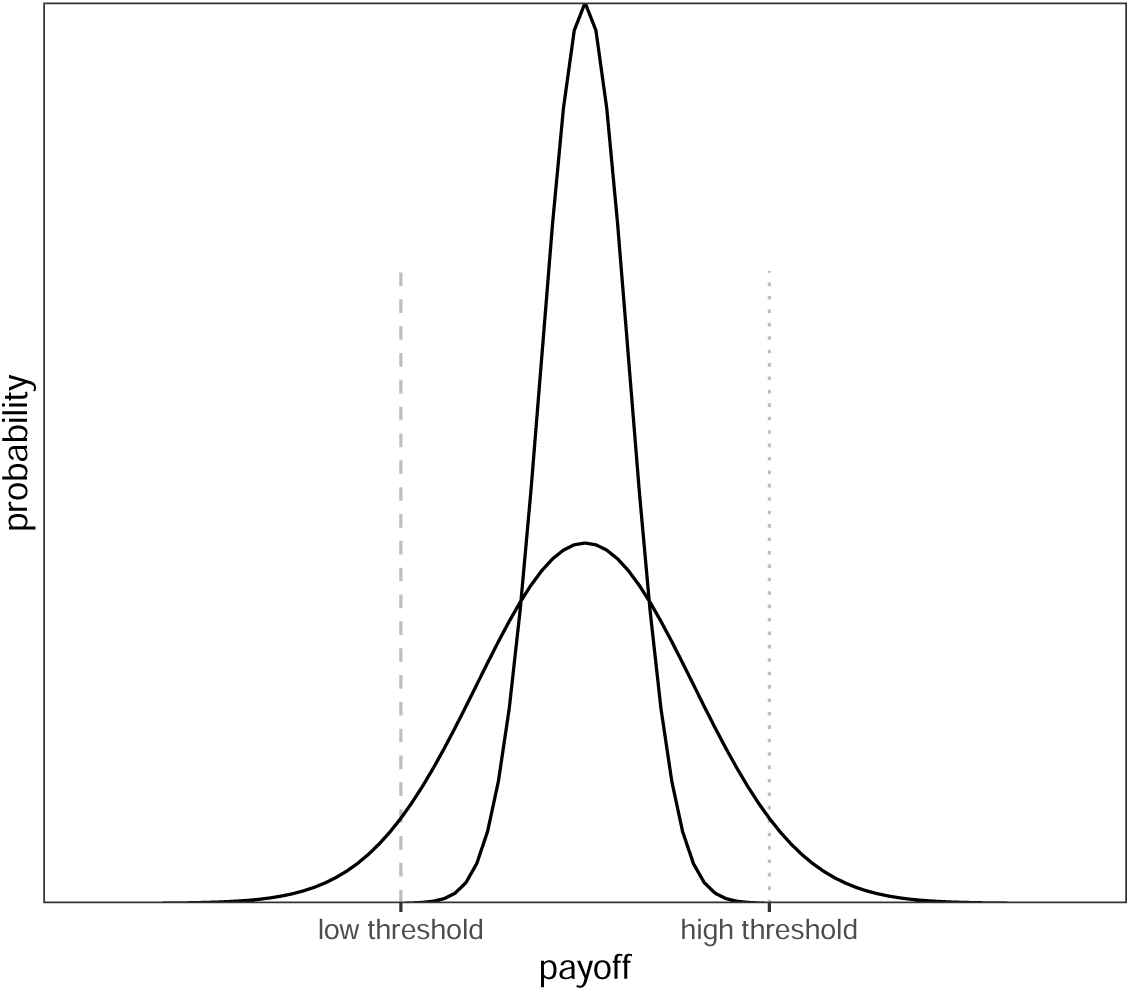
Survival thresholds. When fitness drops to zero below the low threshold (dashed line), the low-risk option (narrow distribution) reliably meets the threshold, favouring risk aversion. When fitness drops to zero below the high threshold (dotted line), only the high-risk option (wide distribution) offers a chance of passing the threshold, favouring risk proneness.

Comparable cutoff points exist in academic careers: Publication records are explicit criteria in decisions such as being awarded a PhD (Altenberger, Leischik, Vollenberg, Ehlers, & Strauss, 2024; Muijrers, 2000), grant funding (Simsek, de Vaan, & van de Rijt, 2024; van den Besselaar & Leydesdorff, 2009), tenure-track hiring (van Dijk, Manor, & Carey, 2014), and promotion (Schimanski & Alperin, 2018). Some of these cutoffs are absolute (e.g., PhD regulations or tenure contracts specifying minimum publication numbers), while others are relative and depend on the pool of competitors (when a job goes to the best candidate, one individual’s success diminishes another’s chances). Such relative thresholds represent *competition*. In the following, survival thresholds (absolute) and competition (relative) are treated as separate concepts.

## Simulation model

We develop an evolutionary agent-based model which simulates a population of researchers who test hypotheses, (attempt to) publish the results either as Registered Reports or as standard reports, accumulate the payoffs for successful publications, and pass their publication strategies on to the next generation of researchers.

### Research phase

Consider a population of *n* = 500 researchers. Each researcher has a fixed publication strategy *s*, the so-called submission threshold. In each round of the research phase, researchers randomly choose a hypothesis to test in a study. Hypotheses are true with prior probability *p*, which is uniformly distributed^4^ between 0 and 1 and known to the researcher. Before testing their chosen hypothesis, a researcher compares the prior *p* of their hypothesis with their publication strategy *s*. When *p < s*, the researcher chooses to play it safe and conducts a Registered Report to test the hypothesis. When *p* ≥ *s*, the researcher chooses to gamble and tests the hypothesis in a regular study which is then submitted as a standard report.

For simplicity, we assume that *p* is an ideal objective prior and that hypothesis tests are free from additional sources of error. Thus, when a researcher tests hypothesis *i*, they obtain a positive result with probability *p_i_* and a negative result with probability 1 − *p_i_*. If the researcher chose to submit a Registered Report, their study is published regardless of the result and the researcher receives a payoff *b_R_*. However, if the researcher chose to submit a standard report, they face strong publication bias: Only positive results are publishable as standard reports and yield a payoff *b_pos_* = 1, whereas negative results are rejected or file-drawered and yield no payoff, *b_neg_* = 0.^5^

For all variations of the model tested here, we assume that the payoff for a Registered Report falls between these bounds, such that *b_neg_ < b_R_ < b_pos_*, for two reasons. First, publication bias makes negative standard reports less valuable than Registered Reports (*b_neg_ < b_R_*): Negative results may not be published at all or require costly resubmissions and revisions in the standard route, whereas Registered Reports are published regardless of the results. Second, positive standard reports remain on average more valuable than Registered Reports (*b_R_ < b_pos_*) because many high-impact journals do not yet offer Registered Reports, because the flexibility of the standard route allows authors to present results in ways that maximise impact, and because the stricter quality criteria of Registered Reports may increase costs and lower net rewards.^6^

This entire research cycle — choosing a hypothesis, choosing a publication route by comparing hypothesis prior *p* to one’s publication strategy *s*, testing the hypothesis, and receiving payoff *b_R_* for a Registered Report or *b_neg_*or *b_pos_* for a positive and negative standard report, respectively — is repeated *m* times.

### Evaluation phase

At the end of the research phase, researchers’ accumulated publication payoffs *b*_1_ + *b*_2_ + *…* + *b_m_* are translated into fitness *f* . Fitness is calculated with a function characterised by exponent *ɛ*, which determines the shape of the function. *ɛ* = 1 yields a linear function, 0 *< ɛ <* 1 yields a concave function with diminishing returns, and *ɛ >* 1 yields a convex function with increasing returns (illustrated in Figure 1):

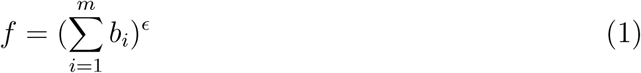

However, two situations may cause a researcher’s fitness to fall to zero even when their accumulated payoffs are non-zero. First, the sum of their payoffs may fall below an absolute survival threshold *δ*, for example when a researcher fails to meet an agreed publication target by the time their ‘tenure clock’ runs out. Thus, when 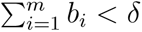, *f* = 0. Second, the sum of their payoffs may fall below a relative threshold *γ*, which reflects the intensity of competition (e.g., for scarce research grants or positions). *γ* is the proportion of researchers who are considered for reproduction. When *γ* = 1, all researchers in the population are considered for reproduction and their fitness is calculated according to Eq. 1. When *γ <* 1, the (1 − *γ*) ∗ 500 least successful researchers receive zero fitness and cannot reproduce. For example, *γ* = 0.1 means that only researchers with accumulated payoffs in the top 10% of the population can reproduce.

### Reproduction phase

Finally, the researchers in the current population retire and a new (non-overlapping) generation of researchers is created. A researcher in the new generation inherits their publication strategy *s* from a researcher in the previous generation with the probability of the previous researcher’s fitness (i.e., the new generation’s publication strategies are sampled with replacement from the previous generation, probability-weighted by fitness). The new generation’s publication strategies are inherited with a small amount of random noise, such that *s_new_* = *s_old_* + *w*, with *w* ∼ *N* (*µ* = 0*, σ* = 0.01). Such hereditary transmission can be interpreted as social learning in which successful researchers are more likely to be imitated (Smaldino & McElreath, 2016; Tiokhin, Yan, & Morgan, 2021), but its primary purpose in the model is technical — it simply provides the machinery for determining which publication strategies are optimal under different conditions.

### Evolved publication strategy

We study how the evolution of researchers’ publication strategies *s* is affected by *b_R_*, *ɛ*, *m*, *δ*, and *γ* (see Table 1). Importantly, *s* is not an absolute decision but determines *how* the choice between formats is made, indicating the amount of risk a researcher is willing to take. Very low values of *s* reflect risk proneness (the researcher nearly always gambles on standard reports, except when hypotheses have extremely low priors), and very high values reflect risk aversion (the researcher nearly always conducts Registered Reports, except when hypotheses have extremely high priors).

**Table 1.** Parameter definitions and values.

| Parameter | Definition | Value [range] |
| --- | --- | --- |
| $n$ | population size | 500 |
| $g$ | number of generations | 250 |
| $p$ | prior probability of hypotheses | uniform [0–1] |
| $b_{neg}$ | payoff for negative standard report | 0 |
| $b_{pos}$ | payoff for positive standard report | 1 |
| $b_R$ | payoff for Registered Report | [.1, .2, . . . , .9] |
| $\epsilon$ | fitness function exponent | [0.2, 1, 5] |
| $m$ | research cycles per generation (‘empirical pace’) | [1, 2, 4, 8, 16, 32] |
| $\delta$ | survival threshold below which fitness = 0, expressed as a proportion of $m$ | [0, .25, .5, .75] |
| $\gamma$ | proportion of most successful researchers selected for reproduction (competition) | [1, .9, .5, .1, .05, .01] |

## Simulation approach

We use the evolutionary mechanism of this agent-based model as a means for identifying optimal behaviour under different conditions. One non-evolutionary alternative is to calculate expected fitness for a wide range of *s* and determine which strategy maximises fitness in each condition. However, this approach cannot easily account for population dynamics and is thus poorly suited for simulating competition effects. We therefore base our study on the evolutionary model, but validate all non-competition analyses against the expected-fitness model and show that both produce virtually identical results (see Appendix).

## Simulation results

We present results in order of increasing model complexity, beginning with simple scenarios to explain the basic functioning of the model before moving to less intuitive results. The analysed parameter values are inherently arbitrary and do not represent real-world units; results are only meaningful in relation to each other.

## Single research cycle per generation, linear fitness function

The first generation of researchers is initialised with randomly distributed publication strategies *s* (uniform [0–1]), which are then allowed to evolve. Figure 3 shows the effect of varying the payoff for Registered Reports *b_R_* with a linear fitness function (*ɛ* = 1), no survival threshold (*δ* = 0), no competition (*γ* = 1), and one research cycle per generation (*m* = 1). Evolved publication strategies *s* approximate the payoff for Registered Reports *b_R_* in each condition (*s_optimal_* = *b_R_*). The reason is that with a linear fitness function and payoffs *b_neg_* = 0, *b_pos_* = 1, the expected fitness from a standard report equals the prior of the tested hypothesis:

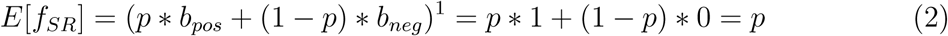

**Figure 3.**
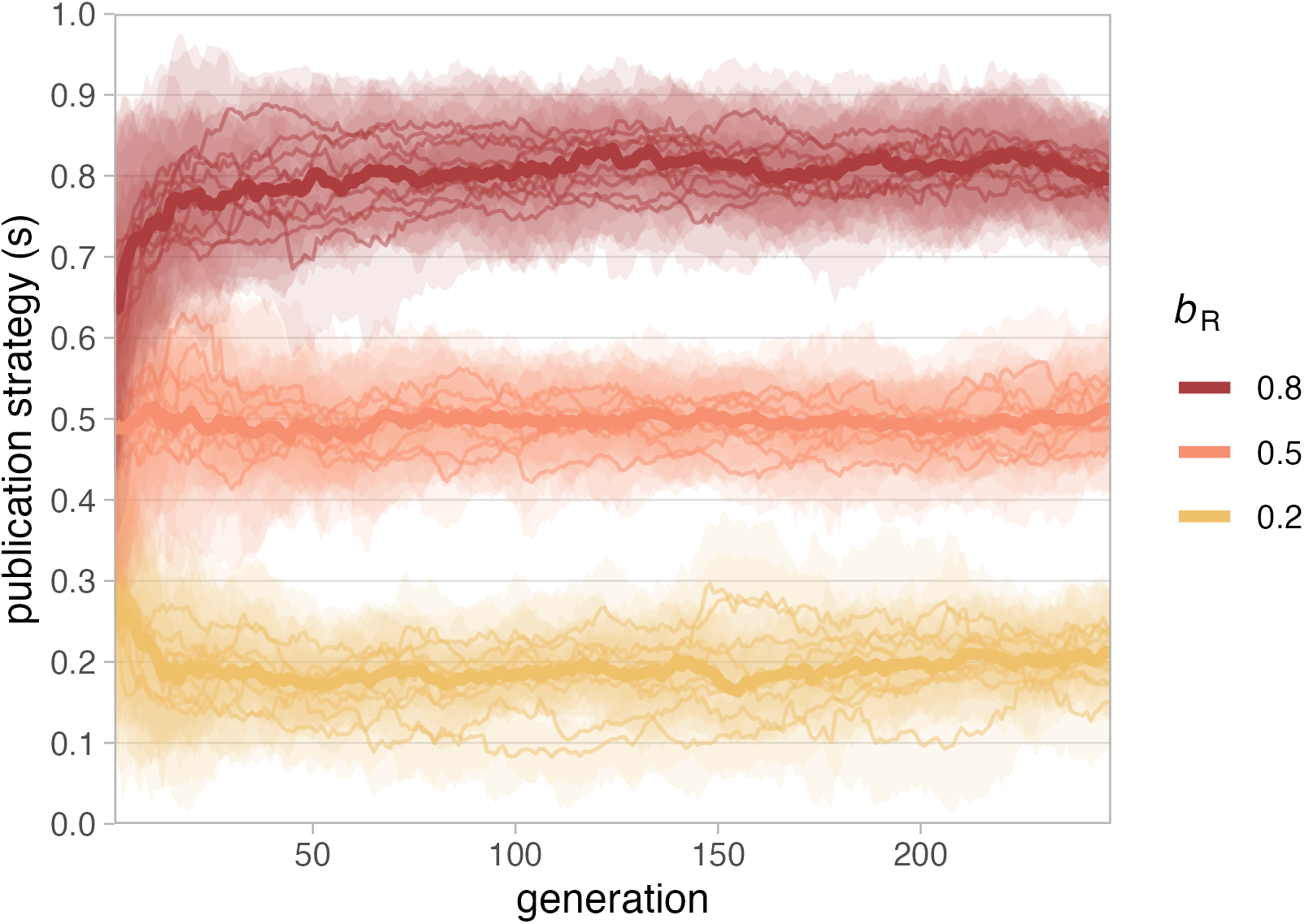
Evolution of publication strategy *s* with 3 different payoffs for Registered Reports (*b_R_*). Parameters: *n* = 500, 250 generations, *b_neg_* = 0 (payoff standard report with negative result), *b_pos_* = 1 (payoff standard report with positive result), *ɛ* = 1 (linear fitness function), *m* = 1 (one research cycle per generation), *δ* = 0 (no survival threshold), *γ* = 1 (no competition), 10 runs per condition. Thin lines: median population *s* per run; shaded areas: inter-quartile range per run; thick lines: median of run medians per condition.

The optimal strategy is therefore to submit a Registered Report whenever the prior of one’s hypothesis is lower than the payoff for a Registered Report (*p < b_R_*). This ensures that researchers get the best of both worlds: For example, when Registered Reports yield the payoff *b_R_* = 0.5, submitting Registered Reports for all hypotheses with a prior *p <* 0.5 protects against losing a bad bet, while gambling on standard reports for hypotheses with *p >* 0.5 maximises winning chances. Every alternative is inferior because researchers who choose Registered Reports for hypotheses with priors higher than the payoff for a Registered Report (i.e., researchers with a strategy *s > b_R_*) forgo winnable gambles and researchers who choose standard reports for hypotheses with priors lower than the payoff for Registered Reports (strategy *s < b_R_*) take unnecessary risks.

## Allowing for non-linear fitness functions

The career benefits researchers receive from publications in the real world are rarely, if ever, linear. Fitness functions may be characterized by increasing returns (i.e., convex), or decreasing returns (i.e., concave). For example, in early career, each addition to a researcher’s short publication record yields increasing returns for their prospects on the job market and their ability to obtain funding (e.g., see Bol, de Vaan, & van de Rijt, 2018). A notable exception may be PhD students who plan to leave academia after their degree, and for whom career returns of publications exceeding the PhD requirements are strongly decreasing (concave fitness function). As researchers gain seniority, the benefits of each additional publication decrease and fitness functions become more concave (Diamond, 1986; Tuckman & Leahey, 1975), although convexity may persist in some cases (e.g., via Matthew effects, Merton, 1968; or competition for priority, Tiokhin et al., 2021).

Figure 4 contrasts the effects of two concave and two convex fitness functions with a linear function for different payoffs for Registered Reports, in the same simple scenario with only one research cycle per generation. Non-linear fitness functions deviate from the linear baseline described above (where the optimal publication strategy is *s_optimal_* = *b_R_*) exactly as expected: When additional payoffs yield diminishing returns (*ɛ <* 1), Registered Reports become more attractive even when they are worth less than the expected payoff for standard reports (risk aversion). Conversely, when additional payoffs yield increasing returns (*ɛ >* 1), Registered Reports are unattractive unless their payoffs are almost as large as those for published standard reports (risk proneness).

**Figure 4.**
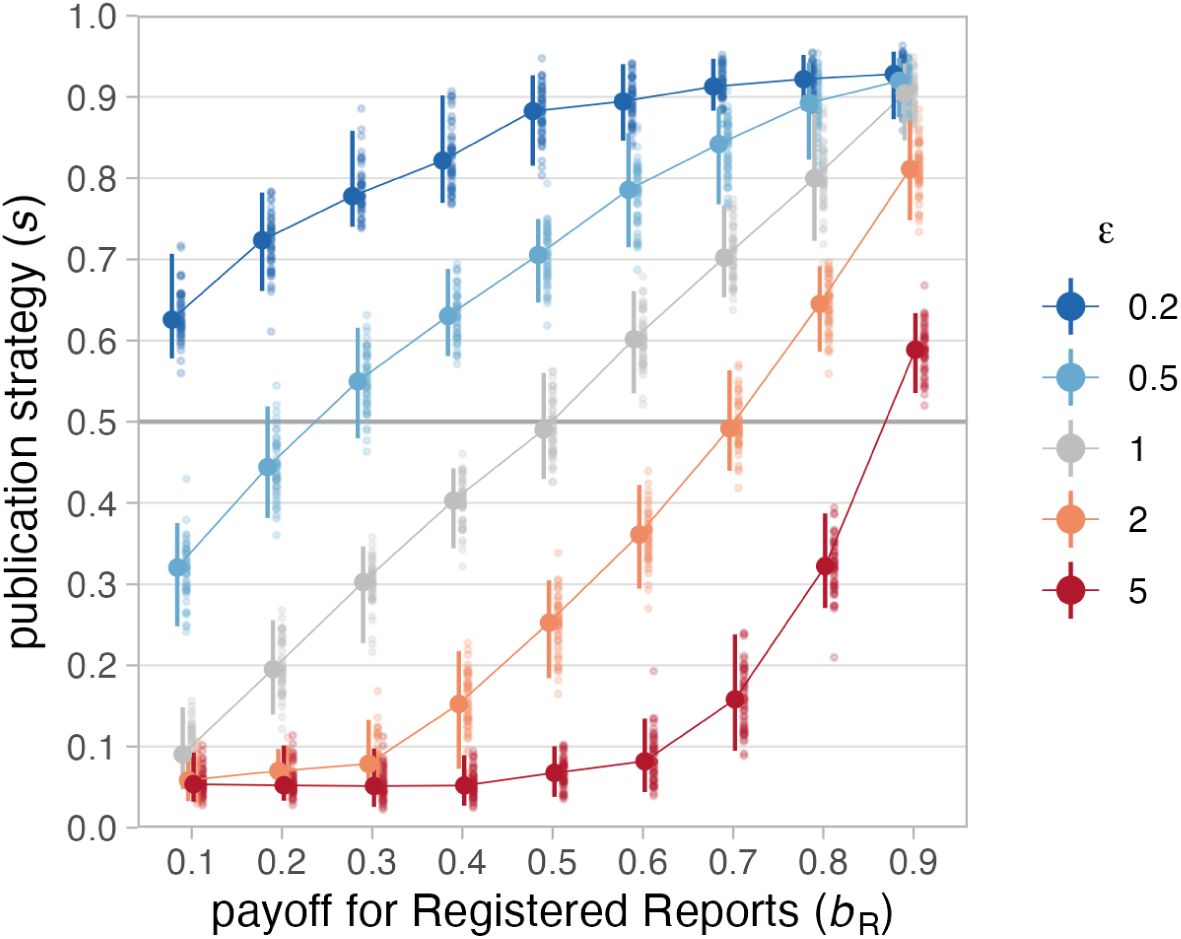
Effect of fitness functions on evolved publication strategies. Median publication strategies after 250 generations (50 runs) for different payoffs for Registered Reports (*b_R_*) and fitness functions (*ɛ*), with one research cycle per generation (*m* = 1), no survival threshold (*δ* = 0), and no competition (*γ* = 1). Blue lines: concave (diminishing returns, *ɛ* = 0.2 and 0.5); red lines: convex (increasing returns, *ɛ* = 2 and 5); grey: linear (*ɛ* = 1). Large dots: median of 50 run medians; error bars: 95% capture probability around median of medians.

When different fitness functions are taken to reflect different career stages, this pattern suggests that Registered Reports should be more attractive for senior researchers and a tough sell for early-career researchers. Yet, limited empirical evidence suggests the opposite: Registered Reports are more likely to have early-career researchers as first authors than standard reports (77% vs 67% in the journal *Cortex*, Chambers & Tzavella, 2022). This could reflect factors not captured in our model, such as younger researchers being more likely to adopt new methods. Other possibilities include early-career researchers intending to leave academia (thus having a concave fitness function), Matthew effects causing convex fitness functions in later career, or that early-career researchers have not yet been subjected to the selection pressures that we model and therefore have strategies that are be poorly adapted to the system.

## Varying the number of research cycles per generation (empirical pace)

Increasing numbers of decision events prior to evaluation may make individuals more risk-prone because single negative outcomes are less catastrophic (Haaland et al., 2019). However, Figure 5 shows that this is not universally true — rather, the effect of empirical pace *m* depends on the shape of the fitness function. Moving up on the y-axis of each panel, the evolved publication strategy *s* decreases (indicating greater risk proneness) only when the fitness function is concave (*ɛ* = 0.2, left panel) but stays constant when it is linear (*ɛ* = 1, middle panel) and even *increases* (indicating risk aversion) when it is convex (*ɛ* = 5, right panel).

**Figure 5.**
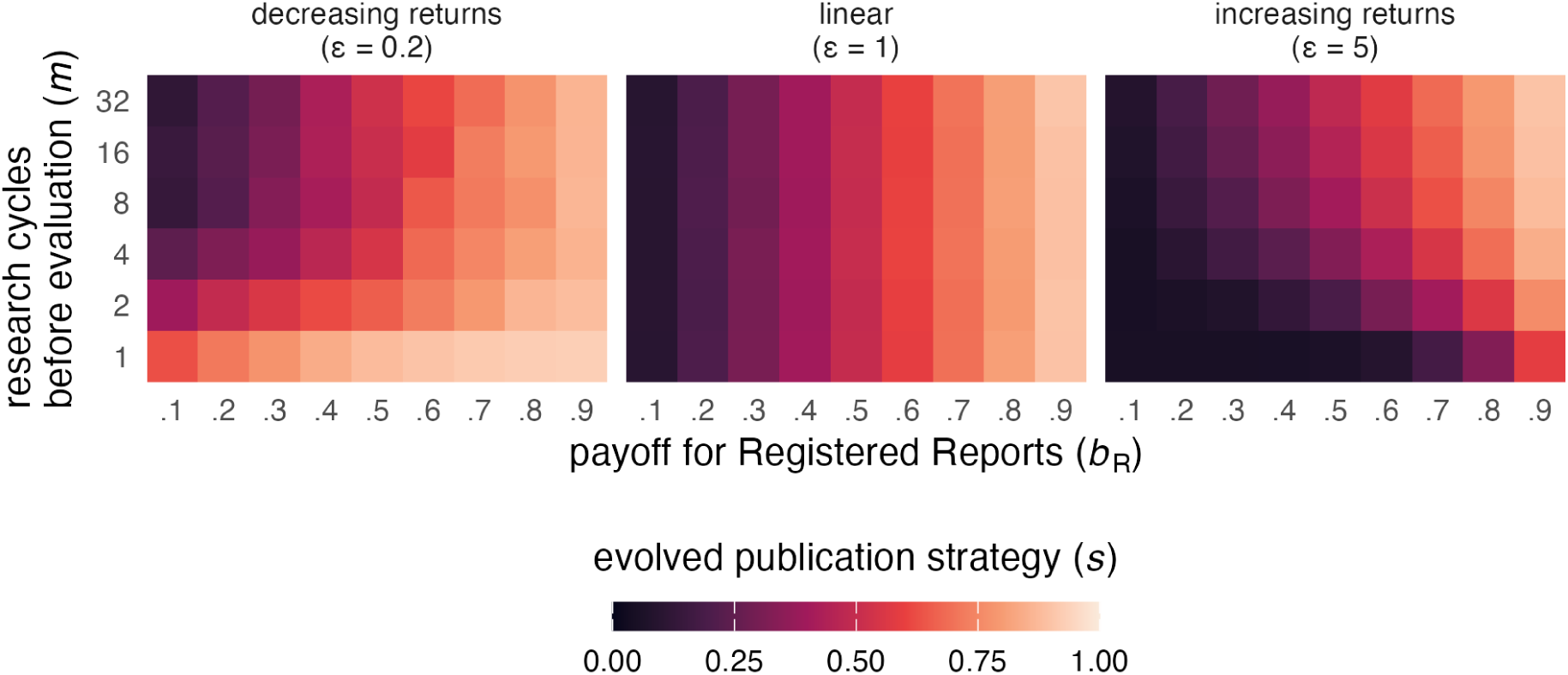
Effect of empirical pace on evolved publication strategies. Tile colour represents median evolved publication strategy *s* of 50 run medians after 250 generations depending on empirical pace (research cycles per generation, *m*), payoff for Registered Reports (*b_R_*), and fitness function (*ɛ*), with no survival threshold (*δ* = 0) and no competition (*γ* = 1).

Why does empirical pace appear to have opposite effects for concave and convex fitness functions? When researchers can complete only one study before being evaluated (*m* = 1, bottom rows), the panels show risk aversion for the concave fitness function

(*ɛ* = 0.2), risk proneness for the convex fitness function (*ɛ* = 5), and a linear strategy for the linear fitness function (*ɛ* = 1). As empirical pace *m* increases, both non-linear panels converge towards the linear case, reflecting the idea that fitness is better captured by the geometric mean when *m* is low and by the arithmetic mean when *m* is high (Haaland et al., 2019).

To illustrate this dynamic, consider two researchers with extreme strategies: Regina Register conducts only Registered Reports (*s* = 1), Darren Daring conducts only standard reports (*s* = 0), with the payoff for Registered Reports *b_R_* = 0.5. With only one research cycle (*m* = 1) and a concave fitness function (*ɛ* = 0.2), Regina’s fixed payoff of 0.5 converts to fitness 0.5^0^.^2^ ≈ 0.87, while Darren’s payoff is 0 or 1 with equal probability (fitness 0 or 1). In a population of 100 Reginas and 100 Darrens, roughly 50 lucky Darrens will narrowly beat all Reginas (1 vs 0.87), while 50 unlucky Darrens are far behind (0 vs 0.87). On average, the Regina strategy is thus more successful. But as the number of research cycles *m* increases, Darrens are no longer split between two extreme outcomes. The law of large numbers causes Darrens’ total payoffs to cluster around the same average as Reginas’, making extreme outcomes rare and optimal strategies converge to the linear case. The same logic applies in reverse when the fitness function is convex (*ɛ* = 5), where Reginas start at a disadvantage (*f* = 0.5^5^ ≈ 0.03) but Darrens’ average payoffs approximate Reginas’ as the number of research cycles *m* increases. Consequently, the top rows (*m* = 32) of the non-linear panels in Figure 5 resemble the linear case.

In terms of academic careers, this pattern indicates that higher empirical pace — e.g., via fast and cheap data collection or greater resources — can cancel out the effects of different career stages. This could partly explain why Registered Reports appear to be less popular among senior researchers (Chambers & Tzavella, 2022) than the effects of diminishing returns alone would predict: academic seniority often comes with resources that boost output per time. Conversely, junior researchers may be especially reluctant to use Registered Reports when they have very limited time or resources before an important selection event, such as on short-term postdoc contracts (R. Müller & de Rijcke, 2017).

## Absolute survival thresholds

The survival thresholds (*δ*) in our model represent absolute publication targets, such as PhD regulations requiring a minimum number of published thesis chapters (Altenberger et al., 2024; Muijrers, 2000) or tenure agreements specifying minimum numbers of publications (Liner & Sewell, 2009; Schimanski & Alperin, 2018).

We investigate thresholds of 25%, 50%, and 75% of the maximum possible payoff per generation. At four research cycles before evaluation (*m* = 4), a survival threshold of *δ* = .75 means that researchers need to accumulate at least 3 points to reproduce. When the threshold exceeds the payoff for Registered Reports (*δ > b_R_*), Registered Reports alone are not enough to ensure reproduction (*b_R_* values to the left of the yellow line in Figure 6). When a Registered Report is worth *b_R_* = .7 in the scenario with *m* = 4 and *δ* = .75, a researcher who conducts four Registered Reports receives only 2.8 points and fails to reproduce. When *b_R_* = .8, the same researcher receives 3.2 and successfully passes their publication strategy on to the next generation. One may thus expect Registered Reports to be popular whenever their payoff is at least equal to the survival threshold (*δ* ≤ *b_R_*) and unpopular otherwise.

**Figure 6.**
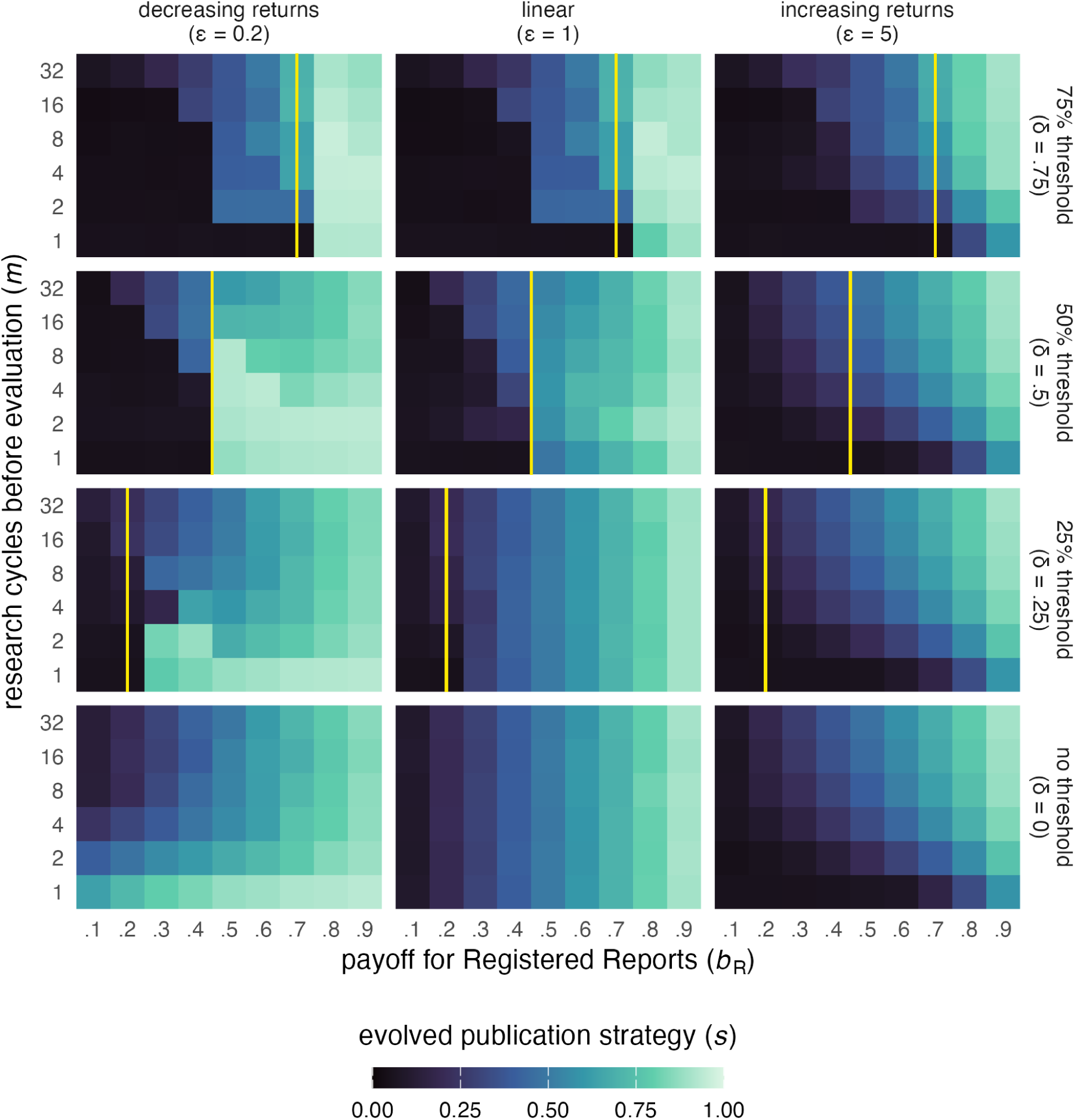
Effect of survival thresholds on evolved publication strategies. Tile colour as in Fig. 5, depending on survival threshold (*δ*, shown as vertical yellow line), fitness function (*ɛ*), empirical pace (*m*), and payoff for Registered Reports (*b_R_*), with no competition (*γ* = 1). Thresholds are set as a proportion of *m*; to reproduce, researchers must accumulate a total payoff exceeding *δ* ∗ *m*.

Figure 6 shows that this holds in many, but not all conditions. Survival thresholds have the largest effect when empirical pace *m* is low and almost no effect when *m* is very high. The effects are stronger for researchers with linear or concave fitness functions (e.g., in advanced career stages). Such researchers almost never choose Registered Reports when their payoff *b_R_* is too low to meet the threshold, especially when they can only complete very few studies before evaluation (low *m*). Conversely, the same researchers prefer Registered Reports more strongly than at baseline when they are worth just enough to pass the threshold, especially when the threshold is high. For researchers with convex fitness functions (*ɛ* = 5), only a high survival threshold of 75% has a visible effect, making Registered Reports even less popular when they have low value (*b_R_ < .*04), especially when empirical pace *m* is low.

In real-world terms, survival thresholds are almost irrelevant when researchers can complete studies at a high pace. Researchers with a convex fitness function (e.g., PhD candidates pursuing an academic career) are only affected by high thresholds, which lead them to choose Registered Reports even less often than normal when their value is low. Researchers with a concave fitness function (e.g., tenure candidates) are highly sensitive to the value of Registered Reports *b_R_*: They virtually never conduct Registered Reports when they are not worth enough to meet the threshold, but strongly prefer them when they are.

## Competition

Competition occurs whenever the demand for academic positions or grant funding exceeds the supply. Figure 7 shows that competition generally leads to an avoidance of Registered Reports, as can be seen by the darkening of the plots when moving up from the bottom row of panels. The only exception to this rule is very low competition: When the top 90% are allowed to reproduce (and only the bottom 10% are rejected, *γ* = .9), Registered Reports become more popular than they are in the absence of competition. This effect is strongest for the concave fitness function (*ɛ* = 0.2), where it holds for almost all payoff settings for Registered Reports (*b_R_*) at very low empirical pace *m* and for high values of *b_R_* at high *m*. When the fitness function is linear or convex, Registered Reports are chosen more often only when both *b_R_* and *m* are high.

**Figure 7.**
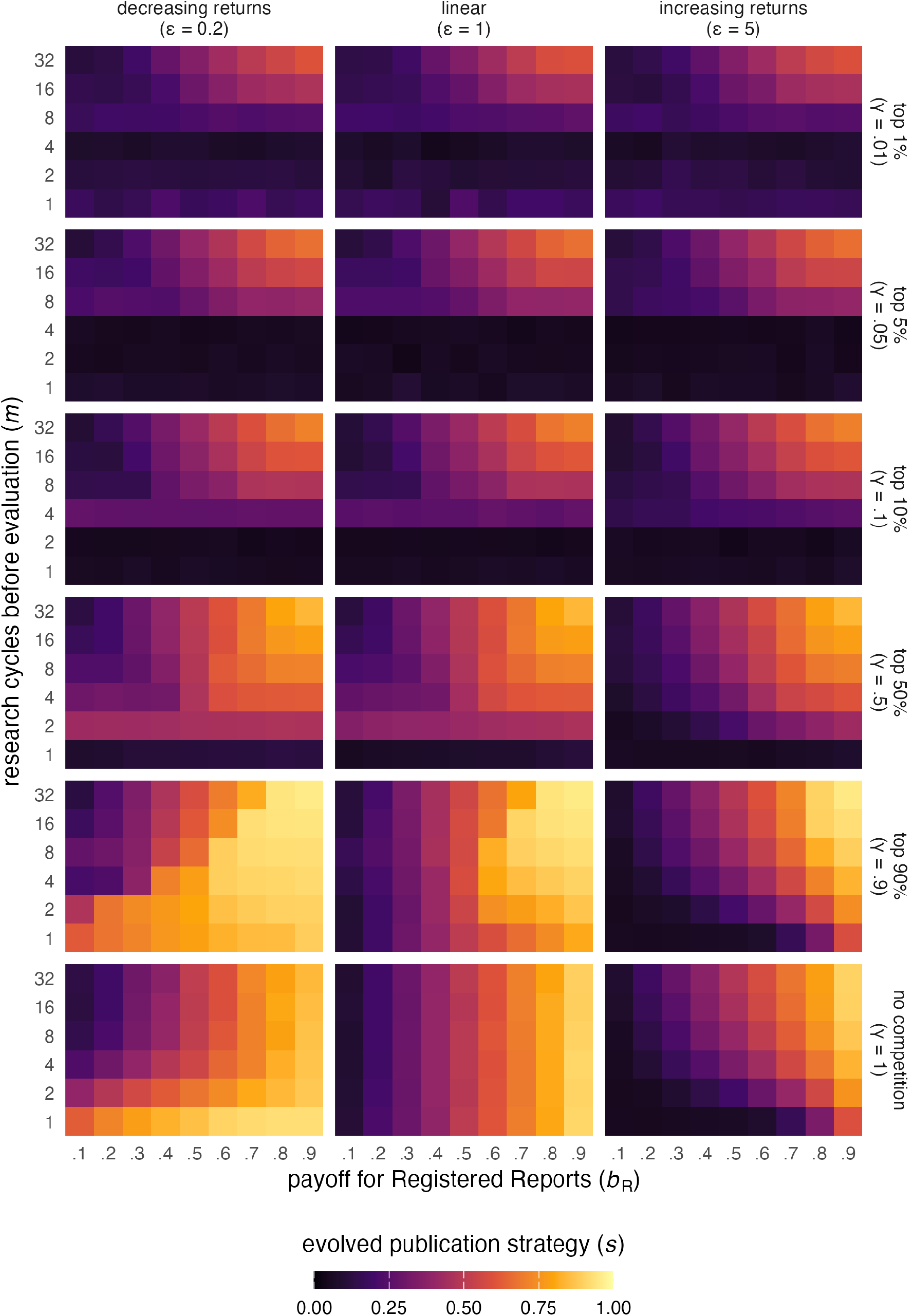
Effect of competition on evolved publication strategies. Tile colour as in Fig. 5, depending on competition intensity (*γ*), empirical pace (*m*), payoff for Registered Reports (*b_R_*), and fitness function (*ɛ*), with no survival threshold (*δ* = 0).

At higher levels of competition (*γ > .*5), the differences between the fitness functions disappear. In all three cases, Registered Reports are essentially wiped out when empirical pace *m* is low, and this effect increases with competition (the higher the competition, the higher *m* must be for Registered Reports to remain viable at all).^7^ Intense competition also negatively affects Registered Reports at high *m*, but here the general pattern of the baseline condition (a linear increase of Registered Reports’ popularity with their payoff *b_R_*) remains intact. This pattern suggests that competition pressures researchers to obtain maximum payoffs. In our model, maximum payoffs are only possible through the combination of choosing standard reports and being lucky (getting positive results). At low empirical pace, this appears to be the best strategy. At higher empirical pace, being lucky in each round becomes so unlikely that choosing Registered Reports at least some of the time becomes a better strategy.

In the academic world, researchers compete for tenured positions and grants, and the intensity of competition varies across fields, countries, grant programmes, etc. Our findings suggest that intense competition may be a significant threat for the viability of Registered Reports, regardless of career stage. This effect is particularly extreme when empirical pace is low. In contrast, very low but non-zero competition *increases* the popularity of Registered Reports, especially when their value is high or the fitness function is concave.

## Discussion

In our model, the standard publication route is a coin toss — the probability of obtaining a publishable result is 50% on average^8^, translating to an expected payoff of 0.5 per study. One might expect payoff-maximising researchers to prefer Registered Reports whenever they are worth more than this expected payoff. This intuition, however, rests on the assumption that career benefits from publications are linear and involve no step changes.^9^ We argue that this assumption is violated in many, if not all, real-world situations. Our results show that when it is, many configurations would lead career-maximising researchers to avoid Registered Reports even when they are worth more than the expected payoff from standard reports.

To summarise, it is useful to take the middle panel of Figure 5 (*ɛ* = 1) as a baseline, where preferences are exactly proportional to the payoff for Registered Reports *b_R_* and unaffected by empirical pace. Compared to this baseline, Registered Reports are *less* popular when a) additional publications yield increasing returns and empirical pace is low, b) researchers face a survival threshold that cannot be met with Registered Reports alone, especially with decreasing returns and low empirical pace, and c) when there is substantial competition (the most extreme effect, causing complete avoidance at low empirical pace). Conversely, Registered Reports are *more* popular when a) returns are decreasing and empirical pace is low, b) their payoff is just high enough to reach a survival threshold and returns are decreasing, especially with low empirical pace, and c) competition is very low but non-zero, especially with decreasing returns or high empirical pace.

Three interaction effects stand out. First, high empirical pace attenuates the effects of all other factors, with high competition as the only exception. Second, the effect of survival thresholds depends strongly on the fitness function, suggesting that publication targets may hit hardest in advanced career stages. Third, the opposite is true for competition, which cancels out fitness-function differences and thus affects all career stages equally.

## Implications

Our model predicts Registered Reports to be least popular when low empirical pace is combined with intense competition or with publication targets that cannot be met with Registered Reports alone. Translated to real-world academia, this suggests that fields or labs in which productivity is limited by lacking resources or the cost or speed of data collection (e.g., research relying on expensive or rare equipment, research on populations that are difficult to access) deserve special attention. Researchers in such situations may avoid Registered Reports when they must achieve publication targets that ask for a minimum number of publications in high-impact journals (e.g., as part of a tenure agreement), or when facing substantial competition for job positions or pressure to obtain competitive grants (e.g., when salary or research time depend on bringing in grants). When competition is high, such researchers may favour standard reports even if Registered Reports are almost as valuable as the best possible outcome from a standard report. Given that the last decades have seen steep increases in PhD student numbers but relatively stable numbers of tenured positions in many countries (Cyranoski, Gilbert, Ledford, Nayar, & Yahia, 2011), substantial competition may in fact be the default in many research fields — which could be one explanation for the currently low market share of Registered Reports.

## Possible interventions to increase the popularity of Registered Reports

Given that Registered Reports reduce bias and allow researchers to publish null results, increasing their prevalence is desirable. What would make Registered Reports more attractive for researchers? Below, we discuss possible intervention targets for each of the factors in our model.

### Relative payoffs

The payoffs associated with Registered Reports may be the easiest factor to intervene on. Net payoffs for Registered Reports can be modified by increasing benefits, lowering costs, or increasing payoff *variance*, each relative to standard reports. Author benefits associated with journal prestige and impact could be raised if more high-impact journals offered the format. Such journals select more strongly on study results (Ceausu et al., 2018; Siontis et al., 2011) and may be reluctant to accept articles before results are known, which could explain why few of them have adopted Registered Reports. It may also explain why *Nature*, the highest-ranking journal offering Registered Reports to date, published only one Registered Report in more than three years after adopting the format (Nature, 2023, 2026). Raising author benefits may thus require convincing more high-impact journals to not only offer Registered Reports, but also to design editorial processes that do not set unrealistic standards.

Reducing the costs of Registered Reports may be a less promising avenue, because much of what makes Registered Reports more costly than standard reports — e.g., higher methodological rigour and larger sample sizes — are ‘good costs’ that increase the quality of the resulting publications (see also Tiokhin et al., 2021). However, journals could preserve quality and still make Registered Reports less costly in *relative* terms by raising the requirements for standard reports (e.g., requiring the same methodological rigour).

Beyond average payoffs, our model indicates that payoff *variance* matters. In the classic Registered Reports model, the pre-data publication guarantee caps not only the worst possible outcome but also the best, because authors must choose a journal before knowing the eventual value of the study. Removing this upper cap could make Registered Reports much more attractive. In April 2021, the platform *Peer Community In* (*PCI*) introduced a journal-independent Registered Reports model in which the review process results in a ‘recommended’ preprint rather than a commitment to a specific journal.^10^ Authors are then free to publish in one of several *PCI* -partnered journals or submit their manuscript to any other journal, which retains the chance of a high-impact publication.

As of July 2026, 41 journals accept *PCI* -recommended Registered Reports without further review. With the growth of this list, and particularly the inclusion of more high-impact journals, the *PCI* model has the potential to change the incentive structure of the format in a profound way.

### Non-linear fitness functions

Our results indicate that Registered Reports may be especially unpopular among researchers with convex fitness functions (accelerating returns, e.g. in Matthew effects). The shape of the fitness function for academic researchers reflects how a field rewards the accumulation of output. Here, bodies responsible for distributing merit, such as hiring and grant committees, wield some influence. Asking candidates to only submit their (e.g.) 5 or 10 best papers for review may help, because additional publications automatically yield decreasing returns. However, if paper quality is at least in part assessed via journal ranking or impact, this method could backfire and make Registered Reports *less* attractive. In fields where competition for priority is common, ‘scoop protection’ policies (Tiokhin et al., 2021) can help dampen Matthew effects that arise from reporting a new discovery first. Another option could be to offer more small research grants in place of fewer large grants to reduce the concentration of merit in the hands of a lucky few (Bol et al., 2018).

### Survival thresholds

Survival thresholds reflect points at which researchers must clear a specific bar to continue their careers. Such bars are often set by faculty committees and can thus be intervened on in principle, the clearest examples being PhD regulations and tenure agreements. One intervention is to change success criteria such that candidates can fulfil them with Registered Reports alone, for example by explicitly offering Registered Reports as a route to tenure or weighting them more heavily than standard reports (for similar calls to emphasise scientific rigour in academic evaluation, see Moher et al., 2018; Schönbrodt et al., 2025). According to our results, such measures could be especially effective in research areas where data collection is slow or costly, and — assuming that very early-career researchers tend to have convex fitness functions that flatten into concave shape over time — that they may be more effective for tenure agreements than for PhD regulations.

### Competition

Our model suggests that very low but non-zero levels of competition may make Registered Reports more attractive, whereas higher levels of competition have a strong negative effect. Reducing the intensity of competition for opportunities such as tenured positions and grants requires additional funding (to increase supply) or restrictions to the applicant pool (to reduce demand), which may be difficult to achieve. Another possibility is to convert traditional grant funding competitions into partial (or ‘modified’) lotteries, where each applicant who clears a minimal quality threshold has the same probability of getting funded (Fang & Casadevall, 2016). Registered Reports should become more attractive if they help researchers build a ‘good enough’ CV to pass the threshold for such partial lotteries. Our results suggest that this can buffer the negative effect of competition on Registered Reports’ popularity in fields with low empirical pace. Reducing competition to a minimum could additionally boost popularity in fields with high empirical pace.

## Limitations and future directions

By design, our model simplifies and exaggerates some aspects of real-world academia and ignores many others. First, we use an extreme, one-dimensional concept of publication bias: all positive results are published, all negative results remain unpublished, and results are determined only by the prior probability of tested hypotheses. Real-world publication decisions depend on many other factors (which may not always favour positive results). Another approach is to model publication bias as favouring results that shift prior beliefs (Gross & Bergstrom, 2021). Compared to this alternative, our model allows a conservative interpretation in which the prior probability of hypotheses simply reflects authors’ predictions of the eventual publication value of different research questions. This interpretation is congruent with Registered Reports and standard reports differing in risk, because publication value depends more strongly on the study results in standard reports than in Registered Reports.

Second, we assume that authors have perfect knowledge of the probability that they will obtain positive (or publishable) results. The assumption that authors have *some* prior knowledge of likely study outcomes is the starting point of our study, because it enables strategic publication behaviour. As long as this assumption holds, adding noise and even bias to authors’ priors would have a diluting effect, but should leave the general simulation pattern intact. More interesting complications arise with individual differences: In our model, researchers who are better at predicting publishability would outperform their peers. In reality, though, certain biases may actually be beneficial, for example if overconfident individuals are also better at convincing editors and reviewers of the value of their work.

Third, researchers in the model cannot choose which hypotheses to test, whereas real researchers routinely choose their own research questions. Researchers who are skilled at identifying hypotheses that are more likely to yield high-impact publications may find Registered Reports less attractive. This would be an example of ability-based risk taking: Individuals with traits or abilities that increase their expected payoff from a risky option should be more risk-prone (Barclay, Mishra, & Sparks, 2018). A more concerning version of this idea is that Registered Reports may be relatively unpopular among researchers inclined to use questionable research practices (or even fraud).

Fourth, we assume that researchers work alone, whereas most real-world publication decisions are made by teams of researchers with different career-related needs. For example, senior researchers may take their PhD students’ needs into account, effectively behaving more in line with a convex fitness function. Real-world strategies can thus be mixtures of the individual strategies in our model. A related consideration is that researchers may compensate for low empirical pace by forming larger teams, sharing payoffs from a limited number of projects. Such behaviour could cause strategies in fields with slow or costly data collection to resemble those expected under higher empirical pace.

Finally, our model ignores publication delays. A common concern is that Registered Reports take longer to publish because authors must await Stage-1 review before collecting data. Standard reports, on the other hand, may have greater delay variance because delays through rejection at multiple journals or reviewer demands for additional studies may create a longer distribution tail. Differences in delay mean or variance could affect researchers’ behaviour, in particular because humans discount delayed rewards (Odum et al., 2020). Researchers who perceive the standard route as faster may prefer it more strongly than our model predicts. Data on the actual distribution of publication delays for both formats, and on researchers’ beliefs about these delays, would be valuable for investigating this possibility.

## Conclusion

Registered Reports are currently the most convincing proposal for curbing publication bias because they ‘[align] what is beneficial for individual scientists with what is beneficial for science’ (Chambers & Tzavella, 2022, p. 29). However, the incentives for choosing the format in the first place may be less well aligned with the interests of scientists. We examined how the pre-data publication guarantee — which reduces the variance of publication outcomes for authors — affects the attractiveness of Registered Reports under different conditions. Our results show that many common features of the academic ecosystem, such as publication targets and competition for tenured positions, promote risk-prone strategies and may lead researchers to avoid the format.

This suggests that the spread of Registered Reports is not simply a matter of time. In psychology, where the format is most popular, a generous estimate puts the annual rate of published Registered Reports at less than 0.1% of the literature^11^ — much lower than the estimated 7% prevalence of preregistration (Hardwicke et al., 2024), even though both reforms were introduced around the same time. Raising their uptake may require additional interventions, such as increasing the value of Registered Reports in hiring and tenure decisions, supporting the journal-independent *PCI Registered Reports* model, or replacing traditional grant competitions with partial lotteries. To the extent that scientific communities have a demand for the kind of low-bias, high-quality evidence that Registered Reports can offer, such measures may be a worthwhile investment.

## Disclosures

### Data, materials, and online resources

Simulated data, code for reproducing all simulations and figures, and the Appendix are available at github.com/amscheel/rr-model. This manuscript was created using RStudio (RStudio 2026.01.1+403, RStudio Team, 2019), R (Version 4.5.2; R Core Team, 2019) and the R-packages *bookdown* (Version 0.46; Xie, 2016), *data.table* (Version 1.18.2.1; Barrett et al., 2024), *ggplot2* (Version 4.0.2; Wickham, 2016), *here* (Version 1.0.2; K. Müller, 2017), *knitr* (Version 1.51; Xie, 2015), *papaja* (Version 0.1.4; Aust & Barth, 2018), *rmarkdown* (Version 2.30; Xie, Allaire, & Grolemund, 2018), *stringr* (Version 1.6.0; Wickham, 2023), and *tinylabels* (Version 0.2.5; Barth, 2022).

### Model development and reporting

The development of the simulation model and the analysis of the simulated data were not preregistered. We report all substantive versions of the model and all parameters that were considered. Parameter values other than those reported here were considered for all parameters except for the prior probability of hypotheses and the payoffs for standard reports and Registered Reports. The values used in the final version of the model were chosen to maximise information value while keeping the number of factor level combinations to a manageable amount. None of the omitted parameter values produced results that contradict the reported results and conclusions. Interested readers are invited to explore alternative settings using the open model code.

### Author Contributions

Conceptualisation: A.S. & L.T., with input from D.L.; data curation, formal analysis, and software: A.S. & L.T.; investigation, methodology, and validation: A.S. & L.T.; supervision: L.T. & D.L.; visualisation and writing — original draft: A.S; writing — review and editing: A.S., L.T., & D.L.

## Acknowledgements

This work was funded by VIDI grant 452-17-013. We thank Julia Rohrer, Wybo Houkes, and Chris Chambers for valuable comments that helped improve this manuscript.

## Footnotes

1 Of course authors may still prefer certain results over others, but the two-stage review process is designed to minimise the influence of such biases on the analysis and interpretation of results.

2 For a complete list of journals currently offering Registered Reports, see https://cos.io/rr.

3 When fitness is evaluated after a single decision event, average fitness across generations is best described by the geometric mean, which is more sensitive to variance than the arithmetic mean because one failure to reproduce can end a lineage. This produces bet-hedging: Risk-averse strategies may be more adaptive across generations even when risk-prone strategies yield better outcomes in any one generation (Haaland et al., 2019). As the number of decision events per generation increases, the arithmetic mean (reflecting additive accumulation of payoffs within a generation) becomes relatively more important, shifting optimal behaviour from risk aversion towards risk proneness.

4 A realistic cross-disciplinary distribution of prior probabilities is hard to establish. For example, molecular epidemiologists may deal with predominantly false hypotheses (Wacholder, Chanock, Garcia-Closas, El Ghormli, & Rothman, 2004), whereas social scientists may often test hypotheses that are trivially true (e.g., due to hidden tautologies, Wallach & Wallach, 1994). The uniform distribution represents a pragmatic, agnostic choice useful for understanding the basic mechanisms at play.

5 Publication bias in the real world is milder and merely implies that negative results are published at a lower rate than positive results. The extreme configuration in our model is a simplification that produces a clearer picture of the general effects of publication bias. Relaxing these settings would produce less extreme results but not change the overall pattern.

6 This paper is predicated on the assumption that most researchers currently view Registered Reports this way — once this changes, the present investigation may happily become redundant.

7 Counterintuitively, the extreme effect of competition at low *m* appears to decrease slightly when competition is highest (*γ* = .01). This is explained by competition becoming so extreme that selection operates more on luck than on strategy: Only individuals who happen to choose hypotheses with high priors (*p*) in every round can reproduce, and the publication strategy *s* becomes relatively unimportant. The relaxed selection on *s* is reflected in greatly increased variance of evolved publication strategies in these conditions (see Figure A4 in the Appendix). This effect of relaxed selection is commonly observed in natural populations (Snyder, Ellner, & Hooker, 2021).

8 Because prior probabilities are uniformly distributed between 0 and 1.

9 Linearity is violated when *ɛ ε*= 1 and also by survival thresholds and competition, which introduce step-changes in the fitness function.

10 https://rr.peercommunityin.org/about^/^

11 We estimate 600,000 journal publications per year in psychology and assume that fewer than 600 are Registered Reports (Chambers & Tzavella, 2022). Total publications estimated via https://lens.org with filters Year Published = 2020--2023, Publication Type = journal article, Field of Study = Psychology.

