## Appendix for "Career incentives make Registered Reports unattractive under realistic academic conditions"

This document contains additional figures for the manuscript '*Career incentives make Registered Reports unattractive under realistic academic conditions*'. This PDF was created in RMarkdown. The reproducible RMarkdown file is available in our online repository at https://github.com/amscheel/rr-model/.

### 1. Alternative simulation model

The results in the main manuscript are based on an evolutionary simulation model. Here, we show that an alternative, non-evolutionary simulation model produces virtually identical results for the effects of non-linear fitness functions ($\epsilon$, Fig. A1), empirical pace ($m$, Fig. A2), and survival thresholds ($\delta$, Fig. A3). This model calculates expected fitness (i.e., the long-run average) for a wide range of $s$ and identifies the publication strategy ($s$) that maximises expected fitness in each condition. Because it is not an agent-based model, it cannot account for population dynamics and therefore cannot simulate the effects of competition ($\gamma$).

The data underlying figures A1–A3 was generated with the alternative expected-fitness model. The simulation code for this model and all figures is available in the online repository.

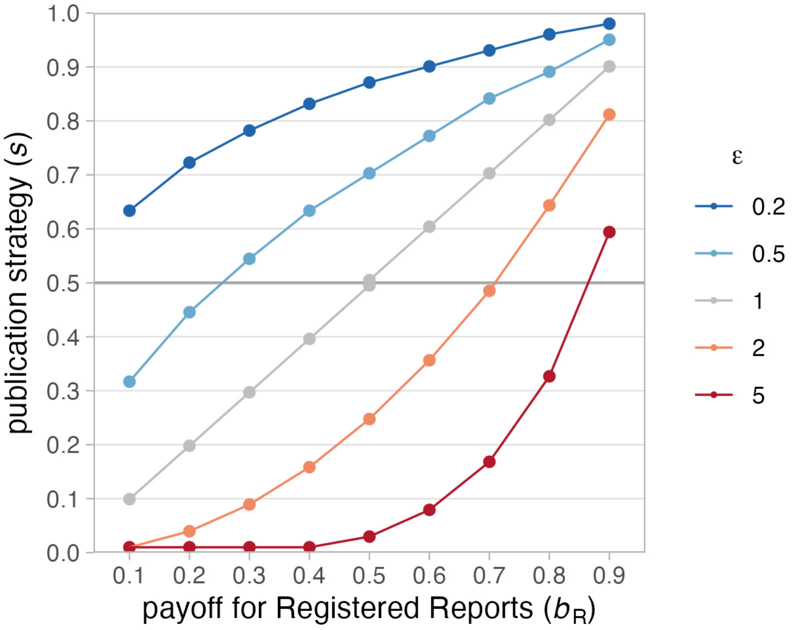


Figure A1: Effect of fitness functions on publication strategies. Shown are publication strategies ($s$) that maximise expected fitness for different fitness functions, with one research cycle per generation ($m = 1$), no survival threshold ($\delta = 0$) and no competition ($\gamma = 1$). Fitness functions with $\epsilon = 0.2$ and $\epsilon = 0.5$ (blue lines) are concave with diminishing returns, functions with $\epsilon = 2$ and $\epsilon = 5$ (red lines) are convex with increasing returns; the function with $\epsilon = 1$ (grey line) is linear.

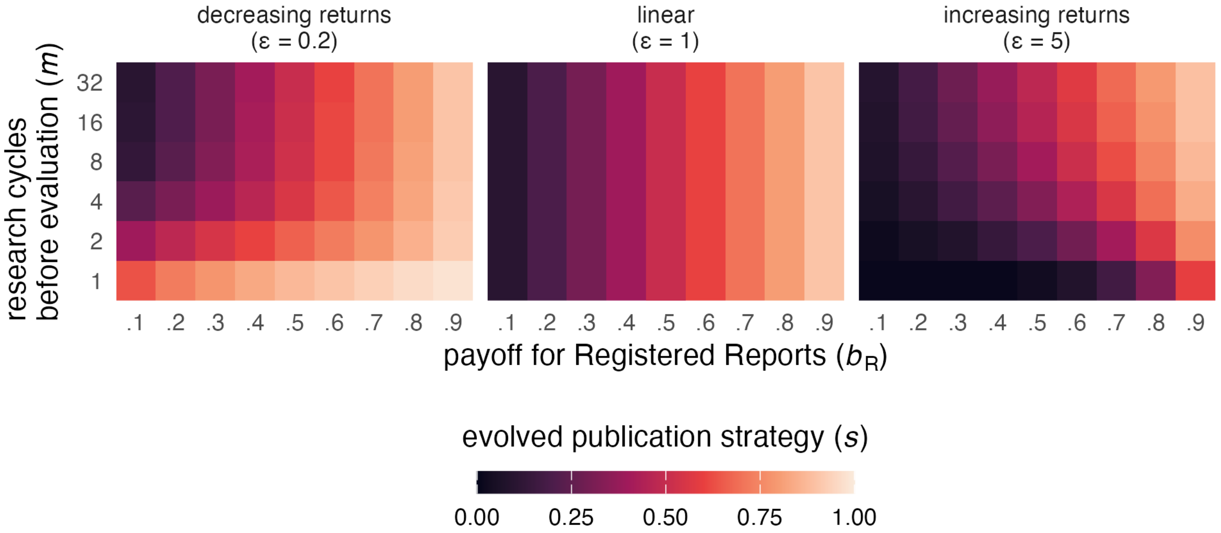


Figure A2: Effect of research cycles per generation (empirical pace) on publication strategies. Shown are publication strategies ($s$) that maximise expected fitness for different fitness functions ($\epsilon$) and numbers of research cycles per generation ($m$), with no survival threshold ($\delta = 0$) and no competition ($\gamma = 1$).

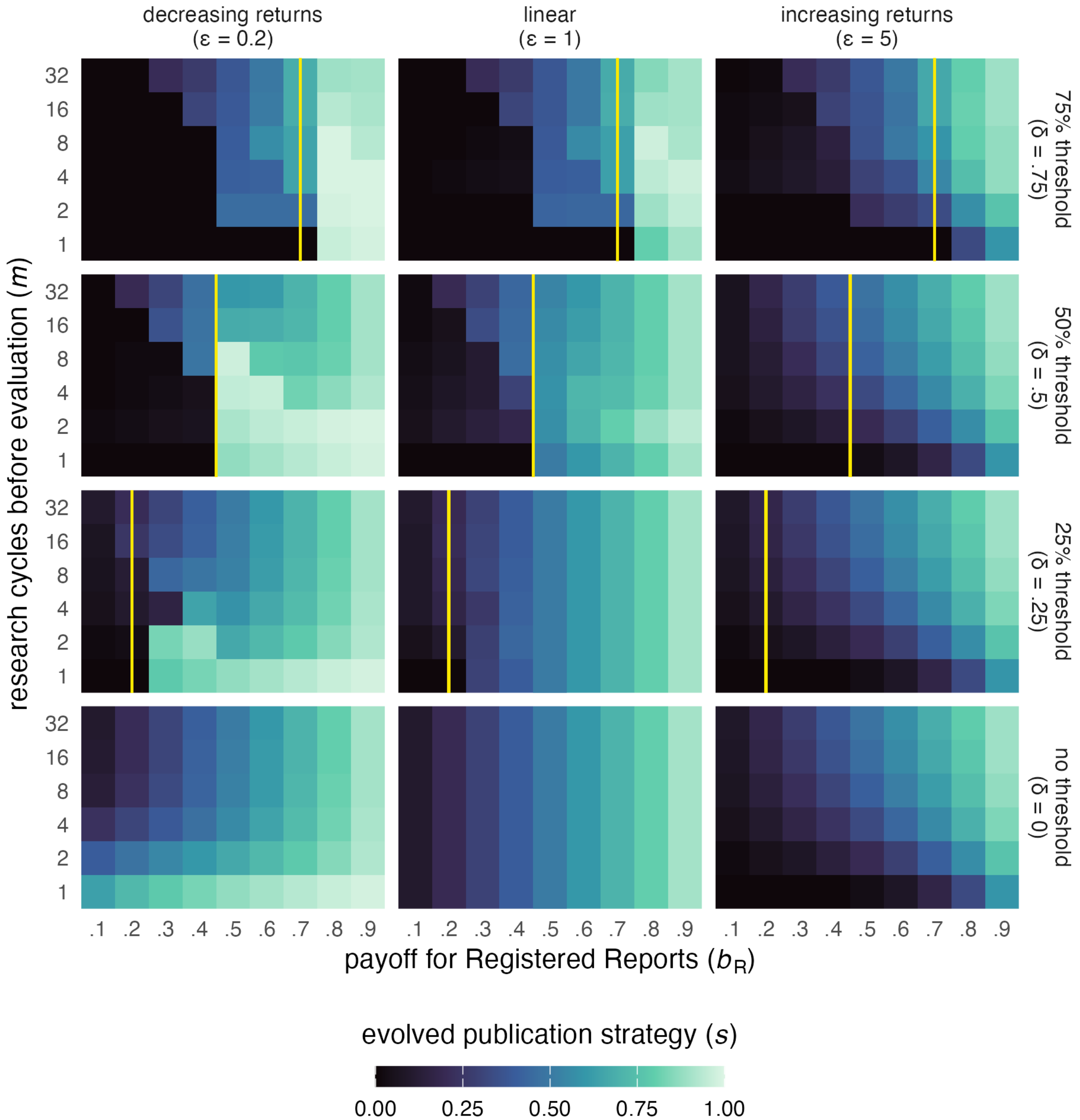


Figure A3: Effect of survival thresholds on publication strategies. Shown are publication strategies ($s$) that maximise expected fitness for different fitness functions ($\epsilon$), numbers of research cycles per generation ($m$), and survival thresholds ($\delta$), with no competition ($\gamma = 1$). Survival thresholds are set as proportions of the maximum possible payoff in each condition and represented by vertical yellow lines.

### 2. Alternative plot for competition results

Footnote 7 in the main manuscript explains that publication strategies paradoxically seem to increase slightly when competition is extremely high and empirical pace is low because natural selection starts operating more strongly on random chance than on heritable traits in this scenario. Figure A4 shows the same data as Figure 7, but plotted in a different way to provide a better insight into this effect: Under low empirical pace, one can clearly see that high levels of competition drastically increase the variance of evolved publication strategies.

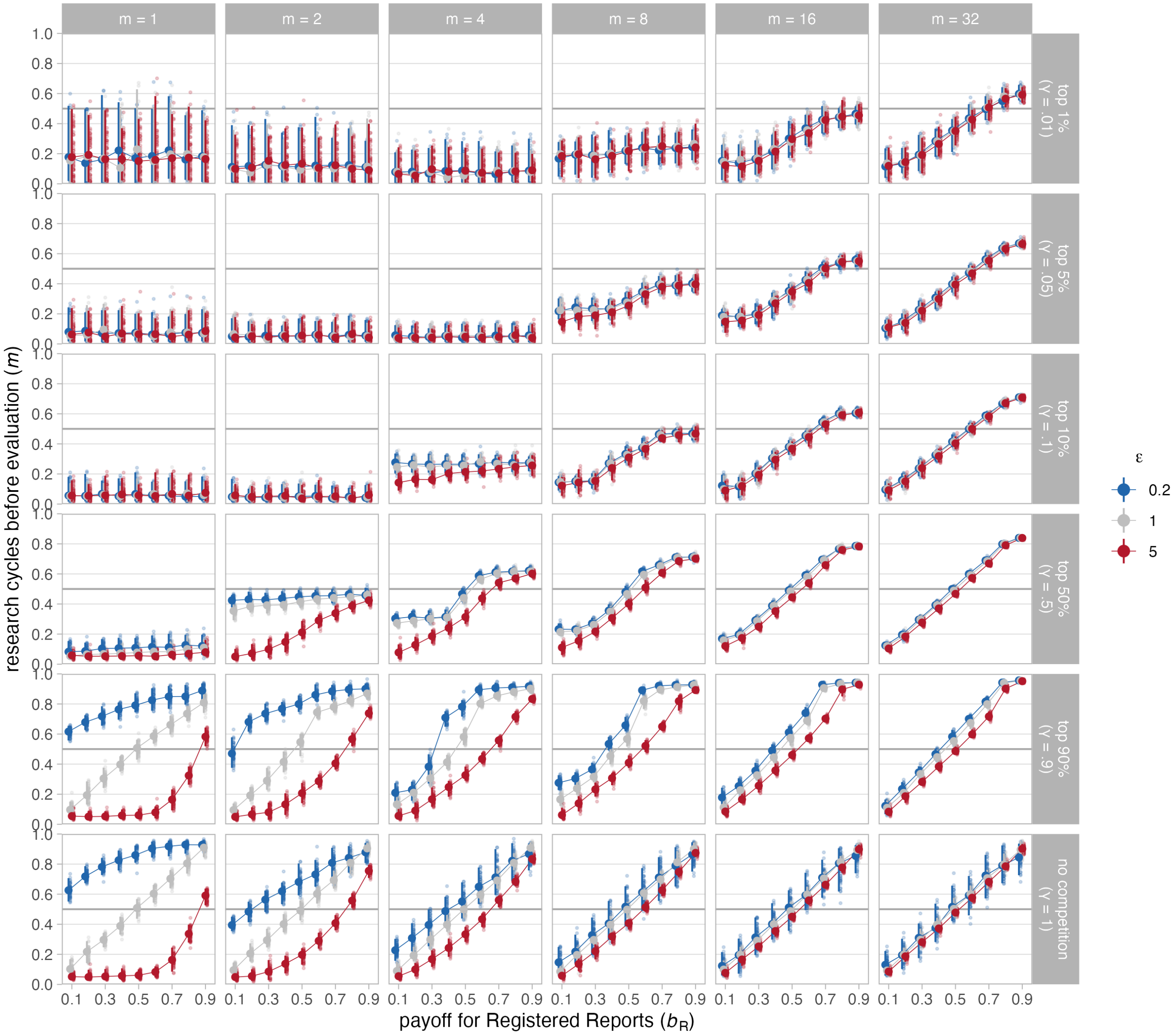


Figure A4: Effect of competition on publication strategies. Shown are median publication strategies in the final ($250^{th}$) generation of 50 runs for different fitness functions ($\epsilon$), numbers of research cycles per generation ($m$), and competition ($\gamma$), with no survival threshold ($\delta = 0$). Small dots represent median $s$ of the final generation in each run, large dots represent the median of these 50 run medians per condition. Error bars represent the 95% capture probability around the median of medians.
